# Co-transfer of functionally interdependent genes contributes to genome mosaicism in lambdoid phages

**DOI:** 10.1101/2022.06.30.498228

**Authors:** Anne Kupczok, Zachary M. Bailey, Dominik Refardt, Carolin C. Wendling

## Abstract

Lambdoid (or Lambda-like) phages, are a group of related temperate phages that can infect *Escherichia coli* and other gut bacteria. A key characteristic of these phages is their mosaic genome structure which served as basis for the “modular genome hypothesis”. Accordingly, lambdoid phages evolve by transferring genomic regions, each of which constitutes a functional unit. Nevertheless, it is unknown which genes are preferentially transferred together and what drives such co-transfer events. Here we aim to characterize genome modularity by studying co-transfer of genes among 95 distantly related lambdoid (pro-)phages. Based on gene content, we observed that the genomes cluster into twelve groups, which are characterized by a highly similar gene content within the groups and highly divergent gene content across groups. Highly similar proteins can occur in genomes of different groups, indicating that they have been transferred. About 26% of homologous protein clusters in the four known operons (i.e., the early left, early right, immunity, and late operon) engage in gene transfer, which affects all operons to a similar extent. We identified pairs of genes that are frequently co-transferred and observed that these pairs tend to be in close proximity to one another on the genome. We find that frequently co-transferred genes are involved in related functions and highlight interesting examples involving structural proteins, the CI repressor and Cro regulator, proteins interacting with DNA, and membrane-interacting proteins. We conclude that epistatic effects, where the functioning of one protein depends on the presence of another, plays an important role in the evolution of the modular structure of these genomes.

**Data summary:** The genomes used in this research are publicly available (Table S1). All supporting data is available in supplementary tables. Source code and documentation to calculate GRR is available under GPLv2 (https://github.com/annecmg/GRRpair).

**Impact statement:** Temperate phages, viruses that can integrate their own genetic material into bacterial genomes, are pervasive mobile genetic elements that can influence bacterial fitness in manifold ways. The *E. coli* phage lambda has been a model phage of molecular biology for decades. Lambdoid phages are highly prevalent in Enterobacteria such as *E. coli* and *Salmonella*, have a mosaic-like genome, the same genome architecture as lambda, and can recombine with phage lambda. Nevertheless, these phages can be very distinct, and no lambdoid core genome exits. Although lambdoid phage genomes have been studied for decades, we know relatively little about how they evolve. Early observations led to the modular genome hypothesis, according to which, phages are assemblages of genetic modules. But what determines the structure of these modules and which genes do preferentially occur together in modules? In this study, we provide answers to these questions using a novel computational approach that allows to infer gene transfer events between distantly related phages despite the absence of a core genome.

We find that co-transfer of functionally related genes is frequent during the evolution of lambdoid phages. This suggests epistatic interactions among these genes, i.e., the co-transferred genes likely need to function together to ensure a viable phage. A prime example is the co-transfer of structural genes, such as genes encoding for the phage capsid or the phage tail. Additionally, we also find co-transfer of known interacting regulatory genes and co-transfer between functionally related genes that have so far been unknown to interact. Together, our analysis provides novel insights into the evolution of temperate phages. Moreover, our approach, which allows to identify gene transfer in the absence of a core phylogeny might be valuable for studying the evolution of other fast-evolving genomes, including viruses of other hosts.

## Introduction

(Bacterio-)phages, i.e., bacterial viruses are considered the most abundant biological entity on earth and play a fundamental role in bacterial ecology and evolution. Phages can either be virulent or temperate. Virulent phages follow the lytic cycle, where phages replicate and lyse their host. Temperate phages can choose between the lytic cycle or a lysogenic cycle during which phage DNA is integrated into the host genome as a prophage that is replicated with the host cell. Prophages, highly relevant for bacterial fitness and evolution (1,2), are very common in bacterial genomes. It is estimated that about 75% of the bacteria are lysogens, i.e., they contain one or more prophages in their genome, which can, in extreme cases, make up to 20% of the bacterial genome content (3–6).

Phages have an astonishing level of genetic diversity (7,8). Even phages that infect a common bacterial host often share no sequence similarity (9–11). This diversity is created and maintained due to a high rate of *de novo* mutations and the acquisition of genetic material through horizontal gene transfer events (12–14). Recent genomic evidence revealed that gene flow is more prevalent in temperate phages, where it likely occurs between infecting phages and resident prophages or between active and defective prophages (15,16). Additionally, gene flow tends to occur among phages enriched for recombinases, transposases, and non-homologous end joining (14). This suggests that homologous and illegitimate recombination contribute to gene transfer in phages (14,17,18).

The “modular theory of prophage evolution” (19,20) states that these extensive gene transfers have created pervasive mosaicism. Accordingly, individual phages are composed of assemblages of shared modules (21–23), where like in a patchwork pattern, almost identical regions can alternate with highly divergent regions. This theory was further supported by several studies suggesting frequent gene flow among these phages (24–30).

Genome mosaicism was first described for lambdoid phages (31), a group of temperate, tailed, double-stranded (ds) DNA enterobacteria phages that share a common genetic architecture and can recombine to form viable hybrids (32). The family of lambdoid phages is sometimes regarded as a single biological species that draws functional genetic modules from a shared gene pool, including phages, prophages, and defective prophages (24–30). Since these modules evolve independently, it has even been suggested that they constitute minimal autonomously functional units, such as groups of genes that must function together or single proteins or even protein domains that function independently (24,33). The genomes of lambdoid phages constitute four different operons: The late operon comprises the morphogenetic proteins, the early left and the early right operon include early transcribed phage genes, and the immunity operon includes the repressor *c*I and the Rex system, known to abort lytic growth of phages (34).

Despite the gene synteny between lambdoid phage genomes, the sequence diversity of lambdoid phages is so high that there exists no lambdoid core genome (26). The genome mosaicism that has been observed in pairwise genome comparisons suggests that some genes are preferentially transferred together. Nevertheless, it is unknown which genes are frequently co-transferred and which functions they encode. This study is based on 26 temperate lambdoid phages with the known ability to perform the lytic and lysogenic cycle (denoted focus phages). To restrict the analysis to active temperate phages, we only included (pro-)phages from databases, with a high genome-wide similarity to the focus phages. Due to the high diversity of lambdoid phages and the absence of a core genome, it is impossible to infer gene transfer with phylogenetic approaches that require a fully resolved phylogeny. Instead, we here build on a previously established two-step approach (14). First, we estimated similarity between two genomes based on protein identities of the homologous protein pairs. Second, we detected gene transfer by identifying highly similar proteins encoded on highly dissimilar genomes. Finally, we inferred co-transfer among lambdoid phages by detecting proteins that are frequently transferred together between the same pairs of genomes.

## Methods

### Genome similarity calculations

For distantly related phages that do not share a core genome, gene repertoire relatedness (GRR) has recently been suggested as a measure to calculate pairwise genome similarity (14). To calculate GRR between a pair of phages, first, homologous proteins are detected as best bidirectional BlastP hits using Blast+ v2.10 (35). Second, global identities (in %) for these homologous protein pairs were determined using powerneedle, a modification of needle from the EMBOSS package for multiple input pairs (36). GRR represents the sum of all global protein identities divided by the number of proteins in the smaller genome and varies between 0 and 100 (in %). The script to calculate GRR for a pair of genomes is available at https://github.com/annecmg/GRRpair.

Average Nucleotide Identity (ANI) was calculated with FastANI using a fragment length of 300 (37).

### Data

#### Focus phages

This study is based on 26 temperate lambdoid phages (Table S1A). Sequences of eight of these phages (HK022, HK97, HK620, Ф80, Gifsy, P22, N15, and λ *PaPa*) were obtained from GenBank. The remaining focus phages were obtained from two sources and subsequently sequenced: Phages with the prefix mEp (Mexican *E. coli* phage) were isolated by Kameyama et al. (38) from human fecal samples in Mexico and were provided to us by Ing-Nang Wang (SUNY Albany, NY). Two phages could not be matched to their original designation and were renamed mEpX1 and mEpX2. A clear plaque mutant of phage mEp043 was used for DNA extraction because its lysate yielded a higher titer. All other phages are likely to be identical to those described in earlier publications (38–40). Phages with the prefix HK were isolated by Elvera and Tarlochan Dhillon in the 1970s and early 1980s from different sources in Hong Kong (some of which are referenced in (41)) and were provided to us by Rodney King (Western Kentucky University, KY).

High-titer lysates of all phages were prepared by confluent lysis (42) and purified (Lambda Midi Kit, Qiagen). DNA content of samples was quantified (Quant-iT DNA assay kit, Invitrogen) and 5 μg of genomic DNA per phage was used to prepare a shotgun library with multiplex identifiers, which was sequenced (Roche 454 Genome Sequencer FLX system with a Titanium kit, Roche Diagnostics). Sequencing was done at the Functional Genomics Center Zürich, Switzerland.

Sequences were assembled using Newbler (Roche Diagnostics) and the assembly software of Geneious 5.4 (Drummond et al. 2011). As coverage was very high, this yielded in all but one case a single contig of the complete genome sequence. We obtained two contigs for phage mEp460, which were then combined using Sanger sequencing. Phage genomes were annotated using RAST searching the Virus domain and using genetic code 11 (Aziz et al. 2008).

The ability to perform both the lytic and lysogenic cycle *in Escherichia coli* is confirmed for the 26 focus phages (Table S1A, (43–46)). The prophages can be induced by mitomycin C and (if they are no clear plaque mutants as indicated with the suffix c-1) are able to integrate into the genome of *E. coli* MG1655 (Fig. S1) (47). We extend this list of 26 focus phages by including (i) temperate phage genomes from NCBI and (ii) predicted prophages from *E. coli*.

#### Temperate phage genomes from NCBI

We searched for virus sequences with host bacteria from NCBI genomes (https://www.ncbi.nlm.nih.gov/genome/browse#!/viruses/) on 16 April 2020. We downloaded the corresponding GenBank files using the Entrez package in biopython. From this list, we retained those genomes that had an ANI of at least 95% to a focus phage resulting in 34 genomes. To discard incomplete prophages, one genome was filtered due to length (length < 27kbp), resulting in 33 phages which we included in the analysis.

#### Predicted prophages from E. coli

We used the search term “Escherichia coli[organism] AND genbank[Filter] AND complete genome[Title] NOT phage[Title]” to find all accession numbers of deposited *Escherichia coli* genomes submitted to NCBI as of 18 May 2020 and downloaded them using the NCBI E-utilities toolkit from the NCBI nucleotide database (48) which resulted in a total of 1115 *E. coli* genomes. Subsequently, we used the downloaded accession numbers on the PHASTER database utilizing their API to find the corresponding PHASTER prophage output of each *E. coli* genome (49). PHASTER predicted a total of 4919 ‘intact’ prophages from these 1115 genomes. Predicted prophage regions were extracted as a FASTA file and annotated using RASTtk (from PATRIC tools) utilizing the Virus domain and genetic code 11 (50). Next, we compared these PHASTER derived prophage sequences to the 26 focus phage sequences using an ANI cutoff of 95% (37). We found 558 prophages with similarity to at least one of the 26 focus phages. Of these, 14 phages were filtered due to length (length < 27kbp or length > 5Mbp), resulting in a final list of 544 prophages.

#### Phage deduplication

The total set of 603 phages (26 focus phages, 33 NCBI phages and 544 *E. coli* prophages) were then deduplicated by grouping highly similar phages by both single linkage and an ANI cutoff of 99.8%. When choosing a representative, we prioritized (i) the focus phages, (ii) the phages from NCBI, and (iii) the phage with the smallest name, i.e., the one originating from the earliest sequencing. We found that one prophage from CP010240.1 contained an insertion element and was thus excluded from the analysis. This resulted in 300 deduplicated phages (26 focus phages, 10 phages from NCBI, 264 prophages).

### Genome groups

We assigned the genomes of NCBI phages and prophages to the same group if they have a GRR of at least 60 to one of the focus phages.

### Protein clusters

First, we performed an all-against-all Blastp with Blast+ v2.10 (35). Significant pairs (e-value <0.001) were aligned with powerneedle and pairs with a global identity of at least 40% were clustered with mcl based on the global identities (51).

### Gene transfer detection

We say that genes from different genome groups were *transferred* if their encoded proteins have a global identity of at least 80%. We calculated PCT (Protein cluster Co-Transfer) between two protein clusters as PCT=GI/(G1+G2-GI), where G1 (G2) is the number of different genome pairs between which protein cluster 1 (2) is transferred and GI is the number of genome pairs between which both clusters are transferred. We define two clusters to be *frequently co-transferred* when their PCT is larger than 0.5.

To compare the PCT statistic to co-occurrence of clusters, we also calculated the PCO (Protein cluster Co-Occurrence) as PCO=PI/(P1+P2-PI), where P1 (P2) is the number of different genomes that contain protein cluster 1 (2) and PI is the number of genome pairs that contain both protein clusters.

### Functional annotation

We manually curated the operon structures by annotating nine model phages based on literature (HK022, HK225, HK620, HK629, Lambda, mEp043-c1, mEp460, P22, Phi80) (52,53). For each annotated protein in a protein cluster, all proteins in that cluster were assigned to the same operon. We observed that the annotation within protein clusters was generally consistent, i.e., cluster members were assigned to the same operon. Only four protein clusters formed exceptions by containing proteins belonging to the early left and to the early right operon; they were not assigned to any operon. Next, we manually curated the operon structure of all genomes by including genes into an operon, if they were in the same orientation and ≤ 10nt from a neighboring annotated gene or if they were in the same orientation, ≤ 200nt from a neighboring annotated gene and the annotation of the other flanking genes does not conflict with the transferred annotation. The resulting annotation is listed in Table S2.

Plotting was done in R using ggplot2 (54), networks were visualized with Cytoscape (55).

## Results and discussion

To identify frequently co-transferred genes and to characterize their function, we estimate gene transfer and gene co-transfer among temperate lambdoid phages (Fig. 1). To distinguish horizontal transfer from vertical inheritance, we aimed to find proteins with high sequence similarity despite low genome-wide similarity of the phage genomes, where they are encoded. Co-transfer is then estimated between protein clusters that are frequently transferred together, i.e., between the same genomes.

**Figure 1:**
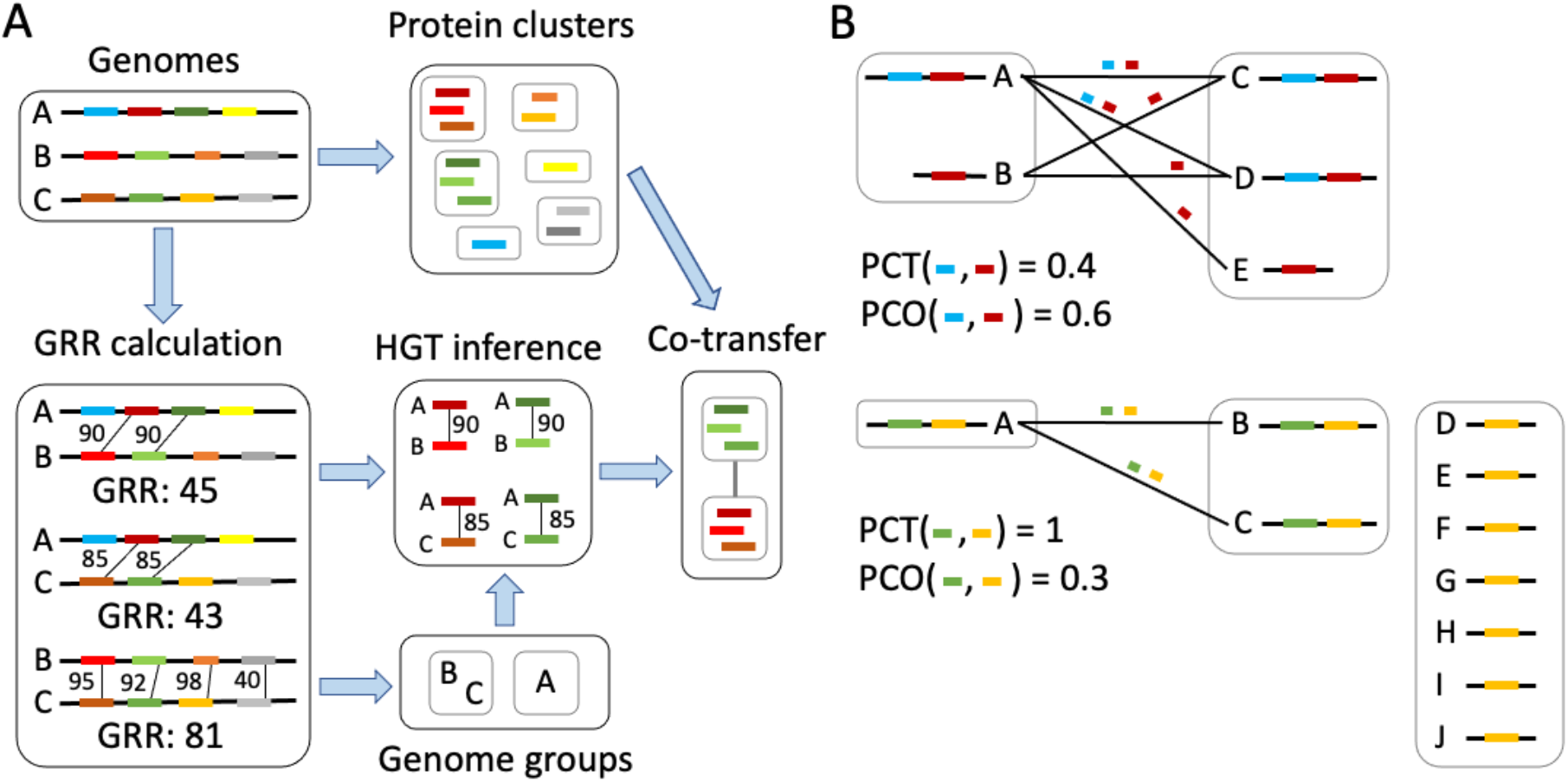
Overview of the approach to detect horizontal transfer and co-transfer. A) For a set of genomes, pairwise GRR is calculated based on protein identities of homologous proteins. Genomes with high GRR are grouped into genome groups. Next, HGT is inferred by detecting highly similar proteins encoded on genomes that belong to different groups. Lastly, co-transfer between protein clusters is inferred when proteins in the cluster are frequently transferred together. B) Example calculations for PCT (protein cluster co-transfer) and PCO (protein cluster co-occurrence). Membership to genome groups is indicated by gray boxes. The blue and red gene are co-occurring frequently (PCO > 0.5) although they are not frequently co-transferred, since PCT < 0.5. In contrast, the green and yellow gene are frequently co-transferred (PCT > 0.5) but do not co-occur frequently (PCO < 0.5).

### Lambdoid phages form distinct groups

To study the genome-wide similarities of lambdoid phages, we analyzed 26 focus phages that were shown to be active in the lab and included 10 related temperate phages from NCBI genomes and 265 *E. coli* prophages after deduplication. We found that 5 NCBI phages and 64 prophages show a gene repertoire relatedness (GRR) of at least 60 to any of the 26 focus phages. This results in a total of 95 phages that were analyzed in the present study (Table S1).

We found that the 95 genomes fall into 12 groups, of which six groups are singletons, two groups contain two genomes, and four groups contain at least four genomes (Fig. 2, Fig. S2A). The groups are characterized by a high GRR (i.e., generally >40) within and a low GRR (i.e., generally < 40) across groups (Fig. 2).

**Figure 2:**
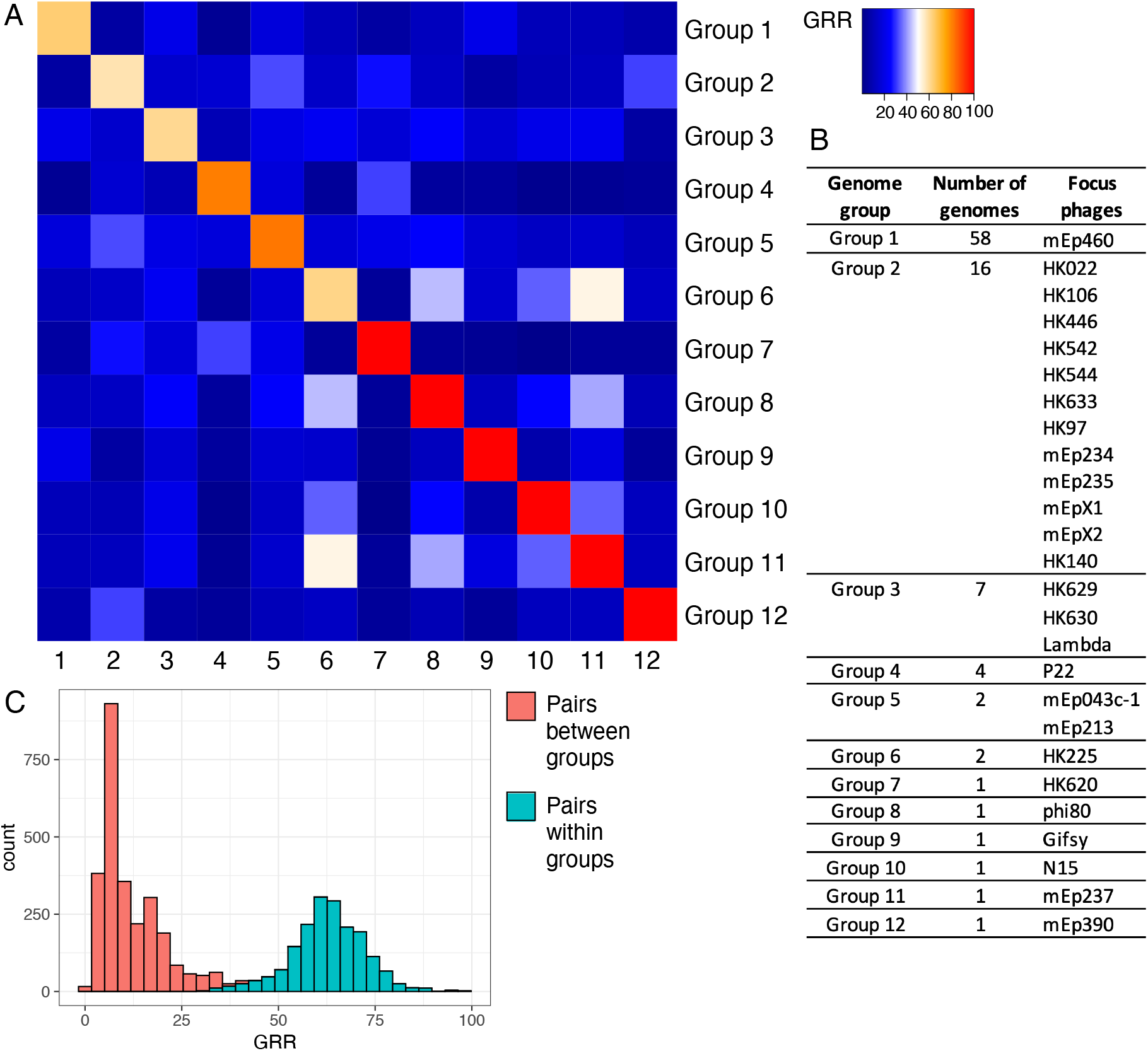
Genome groups. A) Average GRR over genome pairs. Individual GRR values are shown in Fig. S2. B) Number of genomes and focus phages per genome group. C) Histogram over GRR between all genome pairs that fall within (green) or between (red) groups.

Here we used a high GRR cut-off of 60, which allowed us to filter out inactive prophages and distantly related phages, where the life cycle is unknown. This is important as it allows us to define clearly separated groups which ensures that we reliably infer genes that are affected by gene transfer. Nevertheless, additional mosaic phage genomes might exist, that could not be reliably grouped with this approach.

### Protein clustering shows that there is no core genome of lambdoid phages

The 6210 proteins from all 95 genomes were clustered into 1145 protein clusters, of which 608 are singletons (Fig. S3A). We confirmed that lambdoid phages do not contain any core genes, i.e., no gene is present in all 95 phages. Instead, the largest protein cluster contains 83 proteins from 83 different phages belonging to eight different genome groups. Furthermore, only 31 clusters occur in at least 48 (50%) of the phage genomes and no protein cluster contains phages from more than eight genome groups.

Next, we assigned proteins to operons, where 640 (56%) protein clusters could be assigned to a particular operon. We found that the larger protein clusters tend to occur in the late operon (Fig. S3B).

Our analysis confirmed the absence of a core genome of lambdoid phages as previously suggested (26). Thus, a core genome-based phylogeny and traditional phylogenetic methods to infer HGT are not feasible. Instead, we present an approach to infer HGT based on genome groups.

### Gene transfer between distantly related lambdoid phage genomes is frequent and affects all operons to a similar extent

Next, we inferred gene transfer events between phages that belong to different genome groups. Since phages from different genome groups are highly dissimilar (GRR < 40%, Fig. 2C), it is very unlikely that they contain highly similar proteins that have been vertically inherited. Thus, to detect gene transfer events among phages, we searched for pairs of highly similar homologous proteins (proteins in the same protein cluster with an identity of at least 80%) that occur in two different genome groups (Fig. S4A, B). In total, 5238 protein pairs met this condition (Fig. S4B, Table S3). We confirmed that these transferred proteins originate from distantly related genomes characterized by low GRR values that are generally below 40% (Fig. S4C). We found at least one gene transfer in each genome group (Table 1). Gene transfer is more prevalent in some genome groups compared to others, e.g., for groups 3, 5, 7, and 11, more than 50% of the protein clusters are involved in transfers (Table 1). The average proportion of transferred genes per genome also varies between genome groups, ranging between 2% (Group 9) and 77% (Group 7). Note that a very high proportion of genes in a genome can be involved in gene transfer. These gene transfers usually involve multiple partner genomes, consistent with the observations that genome-wide similarities as estimated by GRR are still low between genomes connected by gene transfer.

**Table 1:**
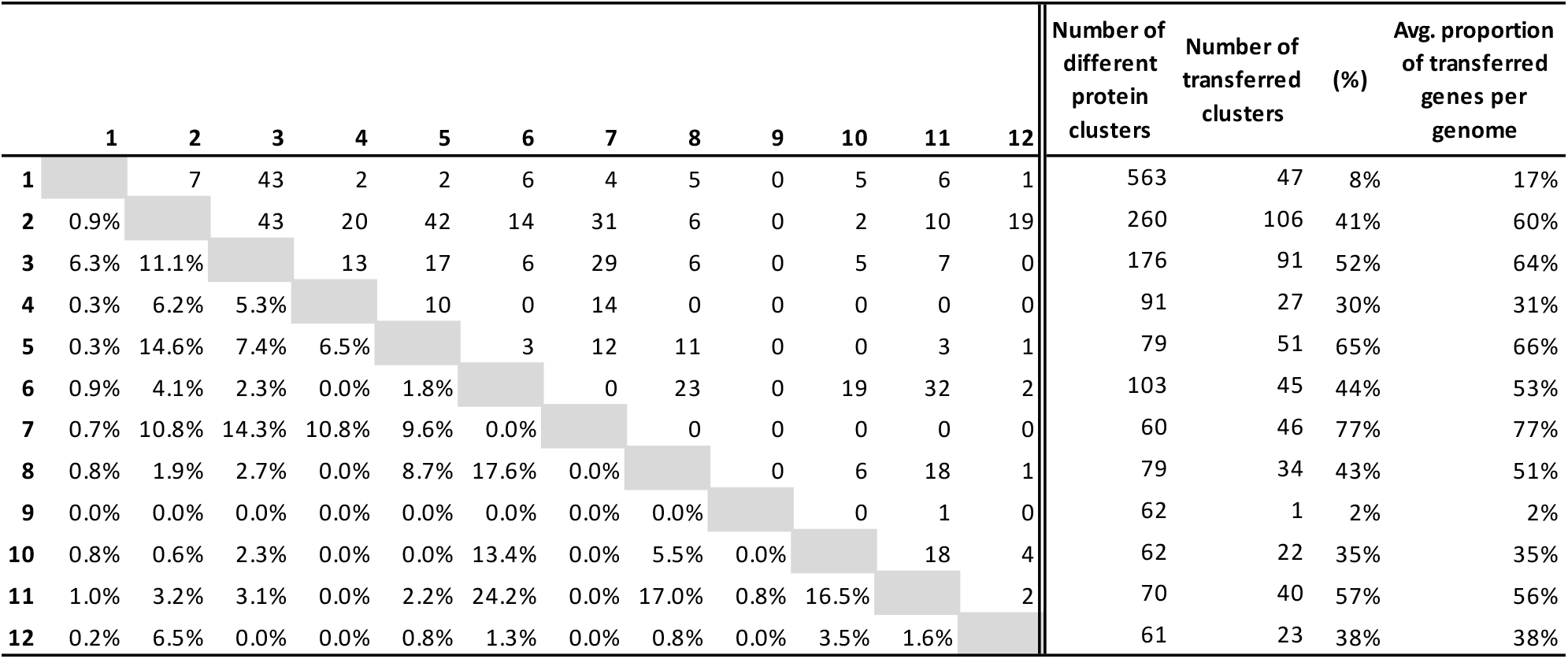
Numbers of transferred proteins between genome groups. Left: Numbers of unique protein clusters that are transferred between each pair of genome groups (upper triangle) and the proportion of these numbers among all different protein clusters that occur in any genome of the two pairs (lower triangle). Right: Total numbers of protein clusters and transferred clusters.

The transferred proteins belong to 180 different protein clusters, where transfers with the focus phages are included in 168 (93%) of these protein clusters. The majority of the 180 transferred clusters fall into known operons, while only 15 (8.3%) were assigned to an unknown operon (Fig. 3A, Table S4). We found that 26% of all protein clusters in operons are involved in gene transfer and that gene transfer affects all operons to a similar extent (Chi-square test, χ^2^=0.038, df=3, p-value>0.05, Fig. 3A, Table S4).

**Figure 3:**
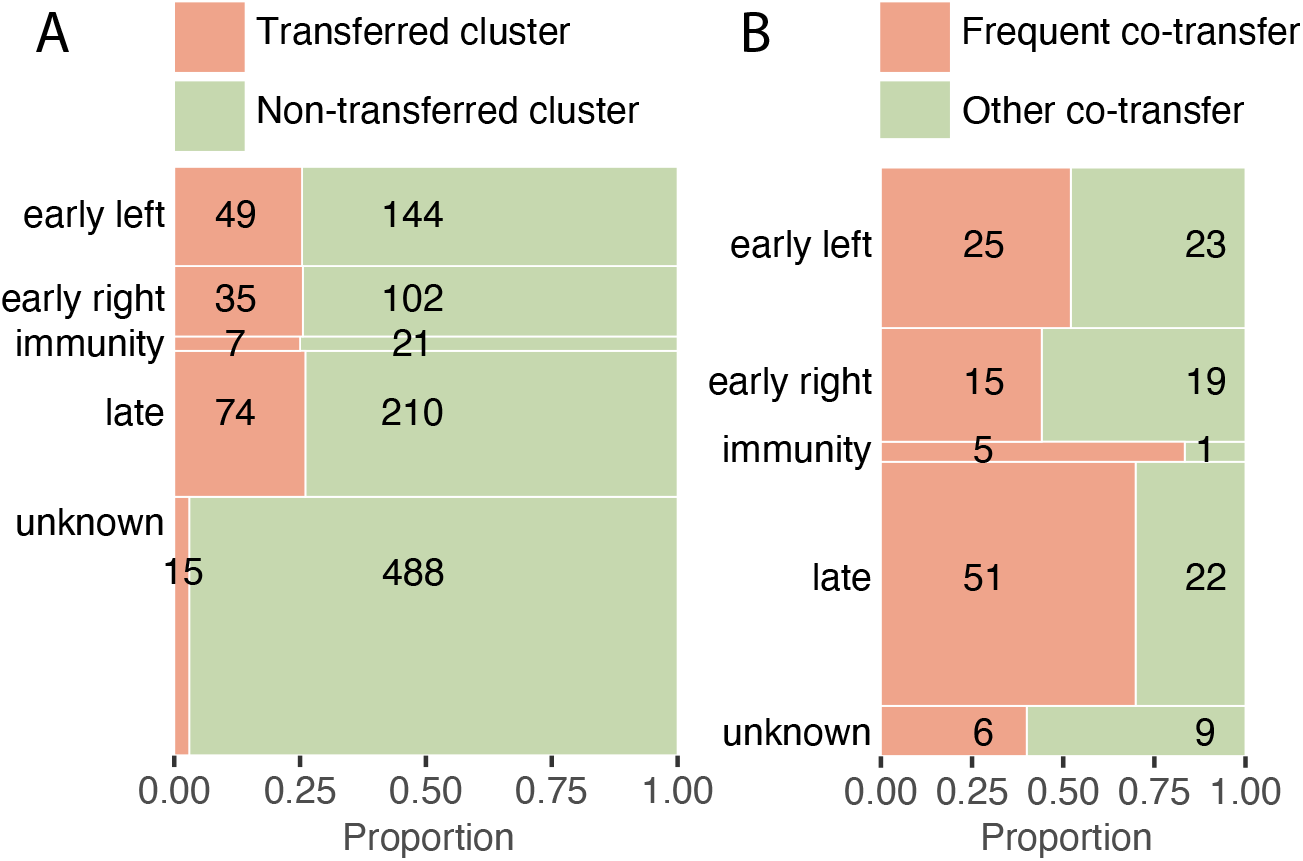
Gene transfers per operon. A) Mosaic plot of protein clusters per operon that are either transferred (orange) or not (green). The proportions between the operons are not statistically different (Chi-square test excluding “unknown”, χ^2^=0.038, df=3, p-value>0.05). B) Mosaic plot of the number of protein clusters that are involved in frequent co-transfers (orange; PCT > 0.5) or other co-transfers (green; PCT ≤ 0.5) per operon. The proportions are significantly different between the operons (Chi-square test excluding “unknown”, χ^2^=9.1, df=3, p-value=0.028). Note that only protein clusters that are involved in at least one co-transfer are included.

We reconstructed HGT among lambdoid phages by identifying pairs of highly similar proteins that are encoded on highly dissimilar genomes. A similar approach has been applied to phages before (14). Here, we extended this approach by focusing on protein clusters instead of protein pairs, which goes beyond previous pairwise approaches and allows us to investigate co-transfer events.

### Co-transfer of genes is particularly frequent in the immunity and late operons

Next, we inferred pairs of protein clusters that are frequently co-transferred (PCT > 0.5), i.e., they are transferred together between more than 50% of the genome pairs where at least one of them is transferred. We found 181 frequently co-transferred pairs of protein clusters (Table S5). These frequent co-transfers involve 102 different protein clusters (58% of all the transferred clusters). Clusters involved in frequent co-transfers occurred more frequently in the immunity and late operon (Chi-square test, χ^2^=9.1, df=3, p=0.028, Fig. 3B, Table S4). The immunity operon is involved in the switch from the lysogenic to the lytic cycle and, in phage lambda, it contains genes encoding for the lysogenic conversion proteins RexA and RexB and the repressor protein cI (53). We find that most of the co-transferred genes encode cI (see Example 2 below), which leads to the overrepresentation of co-transferred clusters in this small operon. The late operon is transcribed late in the phage lytic cycle and it encodes the structural proteins that make up the phage virion. These structural proteins need to successfully form complexes in virion assembly, suggesting that only specific combinations of proteins work well together (53,56), which explains the frequent co-transfer of these genes.

Despite the even distribution of HGT across the different operons, we find an uneven distribution of co-transfer events across operons. The higher occurrence of co-transfer in the immunity and late operons suggests that particular genes in these operons are preferentially transferred together with other genes.

### Frequently co-transferred genes fall into modules of consecutive genes on the genome

To analyze if co-transferred genes might potentially be transferred together by one horizontal transfer event of a DNA segment with multiple genes, we calculated if they are physically close to each other on the genome. We found that the proportion of frequent co-transfer events decreases with physical distance (Fig. 4). The proportion of co-transfers is particularly high for pairs that are directly next to each other or that have at most five genes between them. The high co-transfer rate for genes up to five genes away from each other on the genomes suggests that long segments with more than two genes can get transferred in one event.

**Figure 4:**
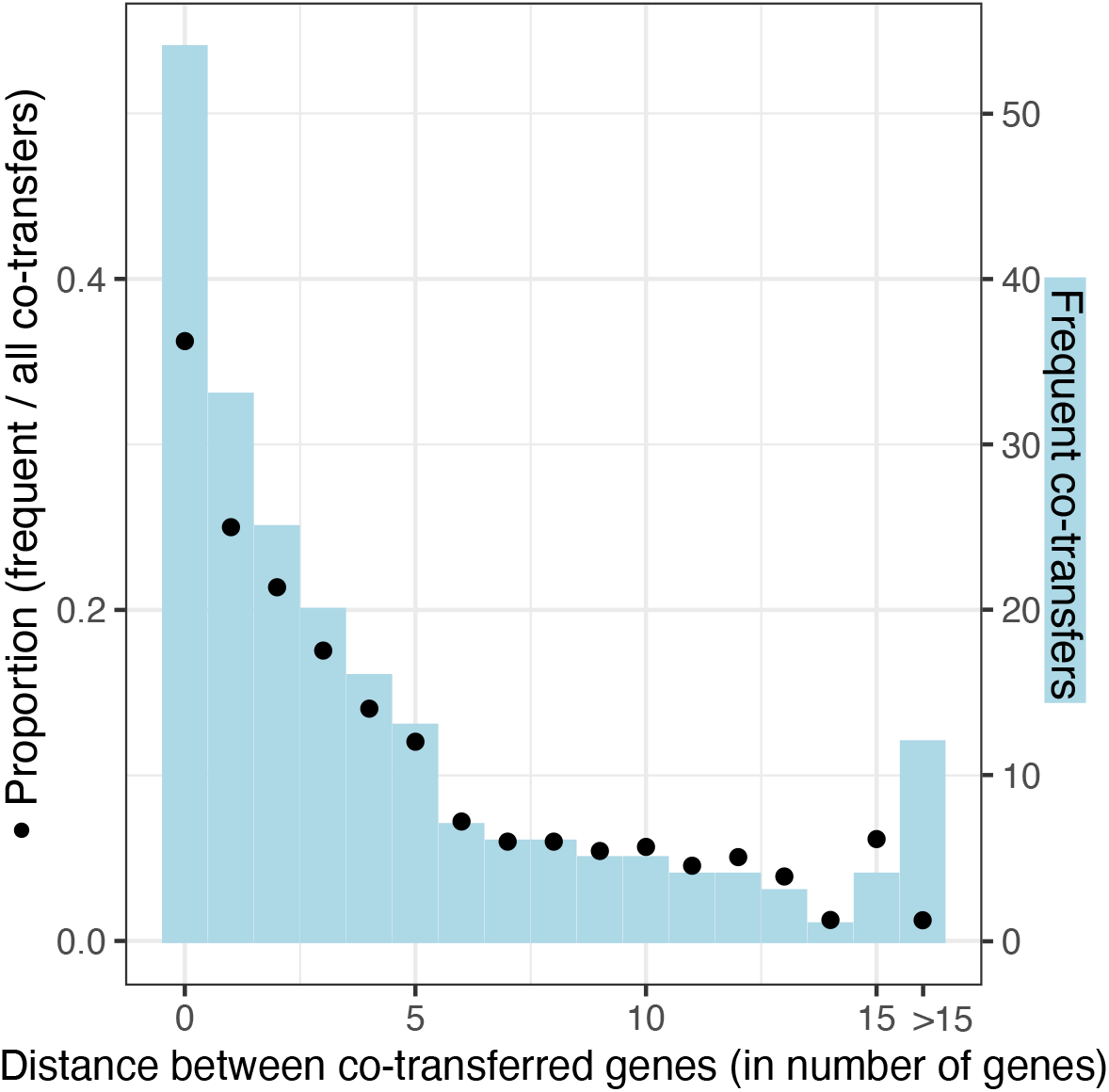
Distance (in number of genes) between co-transferred pairs. The number of frequent co-transfers (PCT > 0.5) per genetic distance is plotted in light blue (right axis) and the proportion of frequent co-transfers among all the co-transfers per genetic distance is shown by dots (left axis).

By linking all 181 pairs of co-transferred protein clusters, we reconstructed a network of 28 distinct modules i.e., connected components (Fig. 5, Table S6). Of these 28 modules, 10 modules contain two protein clusters, seven modules contain three protein clusters, and 11 modules contain more than three protein clusters. In many cases, the co-transferred genes are adjacent on the genome, i.e., the distance in number of genes between them is zero (Table S6). Thus, these modules of consecutive genes lead to small pairwise distances between all pairs of co-transferred genes (Fig. 4). Here we find modules of co-transferred genes. Note that the term co-transfer suggests that these genes have been transferred together in one transfer event, where one phage recombinant is formed by taking up both genes either in one stretch of DNA or in multiple recombination events that occurred in the same infection cycle. Nevertheless, our approach could also pick up sequential transfers, where genes have been transferred after each other during subsequent infections if the respective intermediate stages were viable. Our results suggest that most co-transfers involve adjacent genes on the genome, which makes it likely that they have indeed been transferred together on a single stretch of DNA. Nevertheless, we also observe co-transfer between distant genes, which might be the result of multiple recombination events.

**Figure 5:**
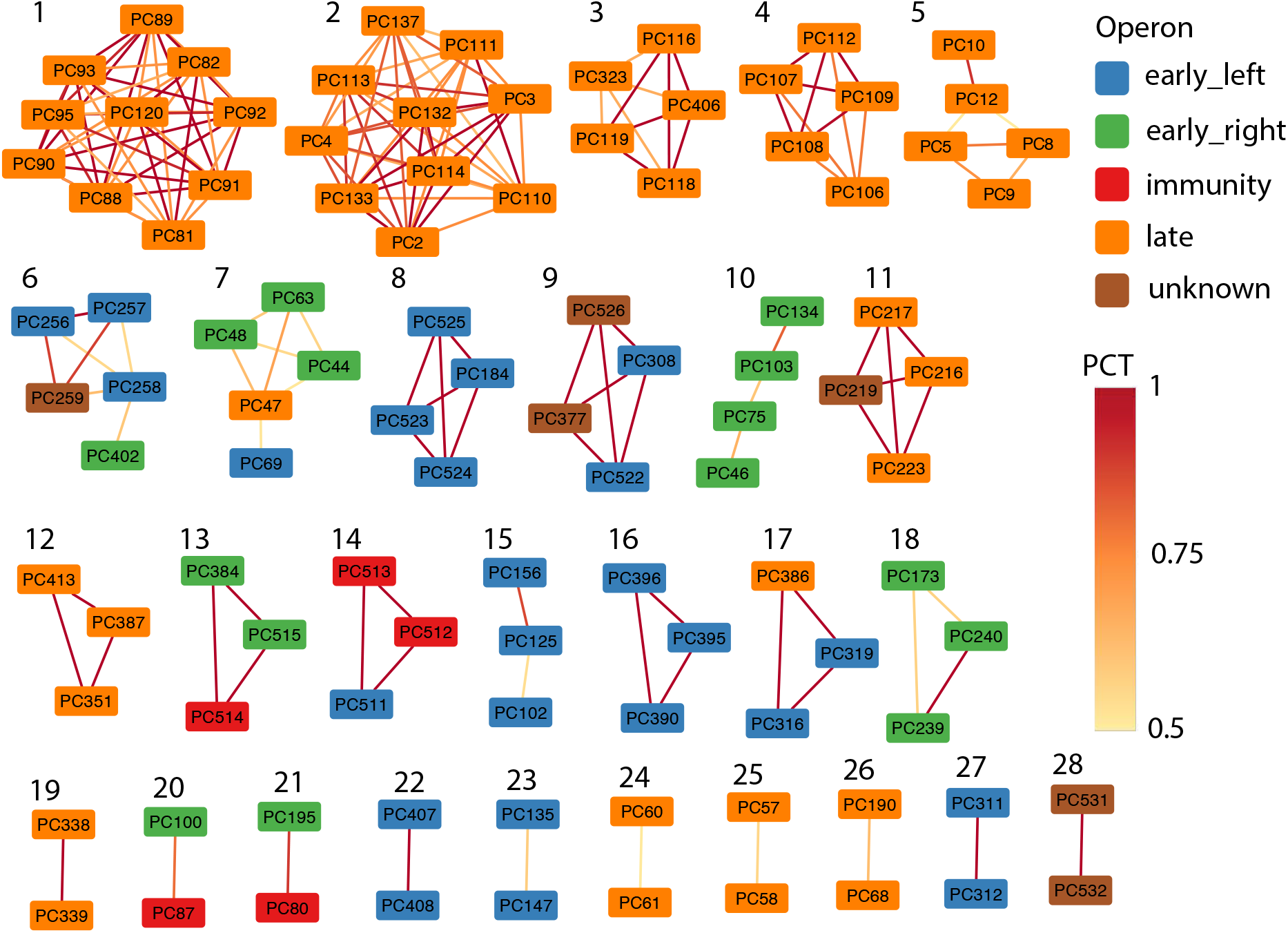
Network of frequently co-transferred clusters. Color-coded by operon. The annotation of the protein clusters is given in Table S6. PCT (protein cluster co-transfer) denotes the proportion how often the genes are transferred together.

### Co-transferred genes are physically closer on the genome than co-occurring genes

In bacteria, co-occurrence of genes across genomes has been used as a signal to detect if genes are functionally associated (57). To compare our measure of co-transfer to co-occurrence, we compared the frequently co-transferred clusters (PCT>0.5) to the frequently co-occurring clusters (PCO>0.5). We observed that co-transfer results in a lower number of pairs compared to co-occurrence (Fig. S5). Furthermore, pairs that are frequently co-transferred have a lower physical distance compared to the frequently co-occurring pairs (Fig. S5).

When comparing co-transfer to co-occurrence, we find that different pairs of proteins are detected with both approaches. With the co-transfer method, fewer genes were found, and these also had a lower physical distance. We thus conclude that the co-transfer method is more specific whereas the co-occurrence method might include more spurious associations. To estimate co-occurrence, we here use a simplistic approach that is based on the number of genomes containing two protein families. Further methods have been developed that need a phylogeny to estimate co-occurrence (e.g., (57)); however, such methods are not applicable to this data set as no core genome exists and thus no core phylogeny can be inferred. The co-transfer approach that we present here circumvents the issue of a missing core phylogeny by detecting co-transferred genes. These gene transfers have been detected as highly similar proteins present on distantly related phage genomes. Instead of estimating a fully resolved core phylogeny, we inferred distinct genome groups and detected gene transfers as highly similar proteins that occur in genomes of different groups. Thus, our co-transfer approach can be applied to genomes where no core genes and thus no core phylogeny exists which makes it particularly very suitable for viruses.

### Frequently co-transferred genes are functionally related

Lastly, we analyzed the functions of co-transferred genes by zooming into the functions of specific modules. We found that the large modules M1 – M5 only involve genes that belong to the late operon. Some modules from the late operon include proteins involved in similar functions suggesting co-transfers of functionally related genes. For instance, module 3 contains only phage tail proteins whereas module 4 contains only phage head proteins. This finding is consistent with the previous observation that head and tail genes belong to separate modules within the genomes of lambdoid phages (52). In the following, we describe three additional examples of co-transfers of functionally related genes.

Example 1: The mEp phages have been selected to contain a variety of immunity groups (38). Thus, our data set contains seven gene clusters of the repressor *cI* (in the immunity operon) and eight gene clusters of the transcriptional regulator *cro* (in the early right operon). Among these, we detected three independent cases of frequent co-transfers that involve cI and Cro (modules 13, 20, 21, Fig. 5, Table S7A). There are some similarities between the proteins in the different clusters. The different Cro proteins that are involved in co-transfer share no sequence similarity, whereas the cI proteins have low pairwise protein identity and were thus clustered separately (Table S7B). Although the co-transfer of this neighboring gene pair can be explained by one transfer event affecting the whole region, it is remarkable that at least three independent co-transfers affected this well-known functional unit, whereas other neighboring genes are not transferred as often. This suggests that different CI and Cro proteins depend on each other.

Example 2: We identified three independent cases of frequent co-transfer that involve the early left protein Kil, an FtsZ inhibitor known to function in host killing (PC184, module 8; PC102, module 15; PC319, module 17, Fig. 5) (58). There is a low pairwise protein identity between PC102 and PC184, whereas the other Kil proteins do not show sequence similarity (Table S7). Interestingly, *kil* is always co-transferred with genes known to encode proteins that are involved in DNA binding (PC527) or in recombination (PC316 – *recT* and PC125 – *erf*). Kil is not essential but becomes critical during conditions of high recombination frequency (58), suggesting that the observed co-transfers are functionally important.

Example 3: Although most modules involve physically close genes, we detected one frequent co-transfer between genes that are distant on the genome (module 5): PC12 encodes a holin protein and is located at the beginning of the late operon whereas PC10 encodes the lambda outer membrane protein Lom and is located at the end of the late operon. Remarkably, both genes interact with the host membrane, where holin is involved in lysis of the inner membrane at the end of the lytic cycle and Lom is integrated in the outer membrane. Lom has so far only been described as a lysogenic conversion protein that belongs to a family of virulence proteins (Pfam accession PF06316) and is involved in adhesion to the eukaryotic host cell (59). Nevertheless, Lom is also strongly expressed during the lytic cycle (60). The co-transfer of *lom* and the holin-encoding gene suggests that Lom might also interact with holin and be involved in bacterial lysis during the lytic cycle.

## Conclusions

Here we show that co-transfer of functionally related genes is frequent during the evolution of lambdoid phages. From this we conclude that the co-transferred genes likely need to function together, suggesting epistatic interactions among gene presences in phage genomes. The importance of epistasis in phages is debated, where recombination has been shown to result in fewer epistatic interactions (61). Nevertheless, the functionally associated frequent co-transfers described here suggest that epistasis is abundant among the frequently recombining temperate phages. We also observe that the co-transferred genes are physically close on the genome. This is expected for recombining organisms, where simulations showed that sexually reproducing organisms evolve modular genomes, where related functions tend to be physically proximal (62–64). Our results confirm this scenario for lambdoid phages, where abundant gene transfer leads to the evolution of a modular genome architecture. Functionally related genes evolved to occur physically proximal on the genomes and gene transfers of functionally related genes maintain epistatic interactions despite frequent gene transfer. Thus, the interplay of epistasis and gene transfer can explain genome mosaicism among phage genomes.

## Authors and contributors

Conceptualization – AK, CCW; Sequencing – DR; Methodology – AK; Formal Analysis – AK, ZB; Data Curation – AK, CCW; Writing – Original Draft Preparation – AK, CCW; Writing – Review and Editing – AK, CCW, ZB, DR

## Acknowledgements

We would like to thank Mumun Gençoglu and Gabriela Klecak for help in the laboratory and Rodney King and Ing-Nang Wang for kindly sharing their phage isolates. We thank Franz Baumdicker and Ing-Nang Wang for valuable discussions on the manuscript.

