## Supplementary figures and table legends for "Co-transfer of functionally interdependent genes contributes to genome mosaicism in lambdoid phages"

### Supplementary material

#### Supplementary figures

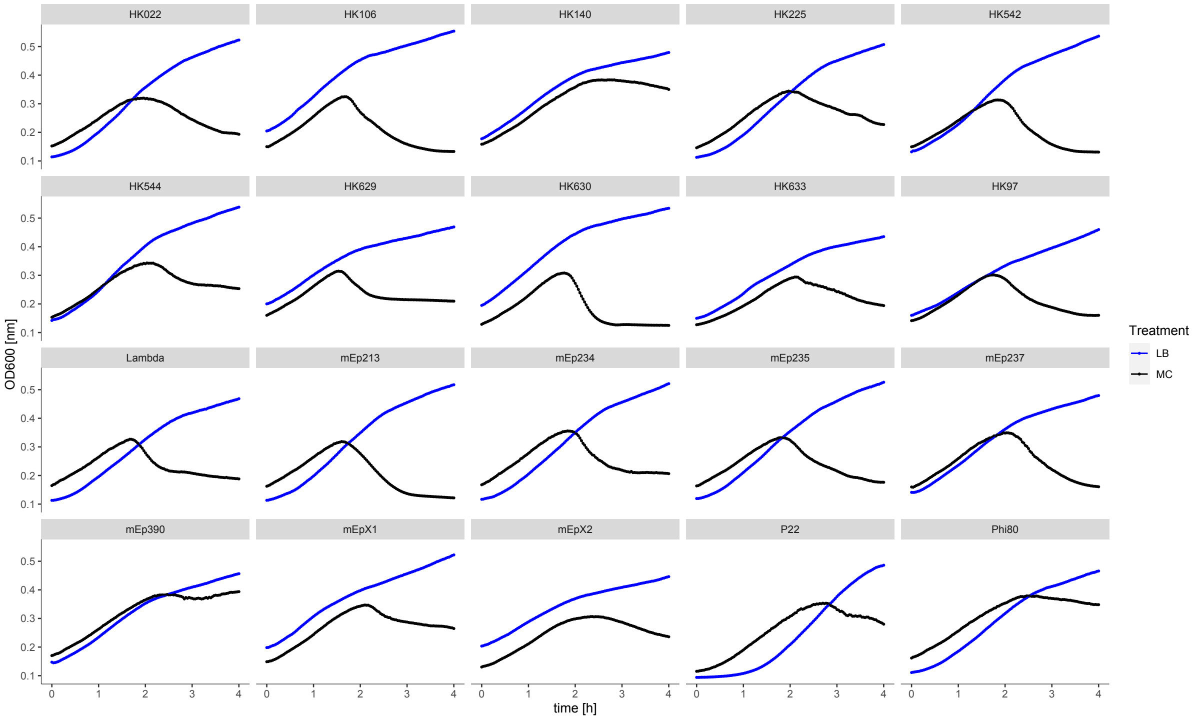


**Figure S1: Induction curves for 20 of the focus phages.** 4-hour growth curves in the absence (black) and presence of mitomycin C (black). The activity of the remaining 5 focus phages has been shown previously (see references in main text).

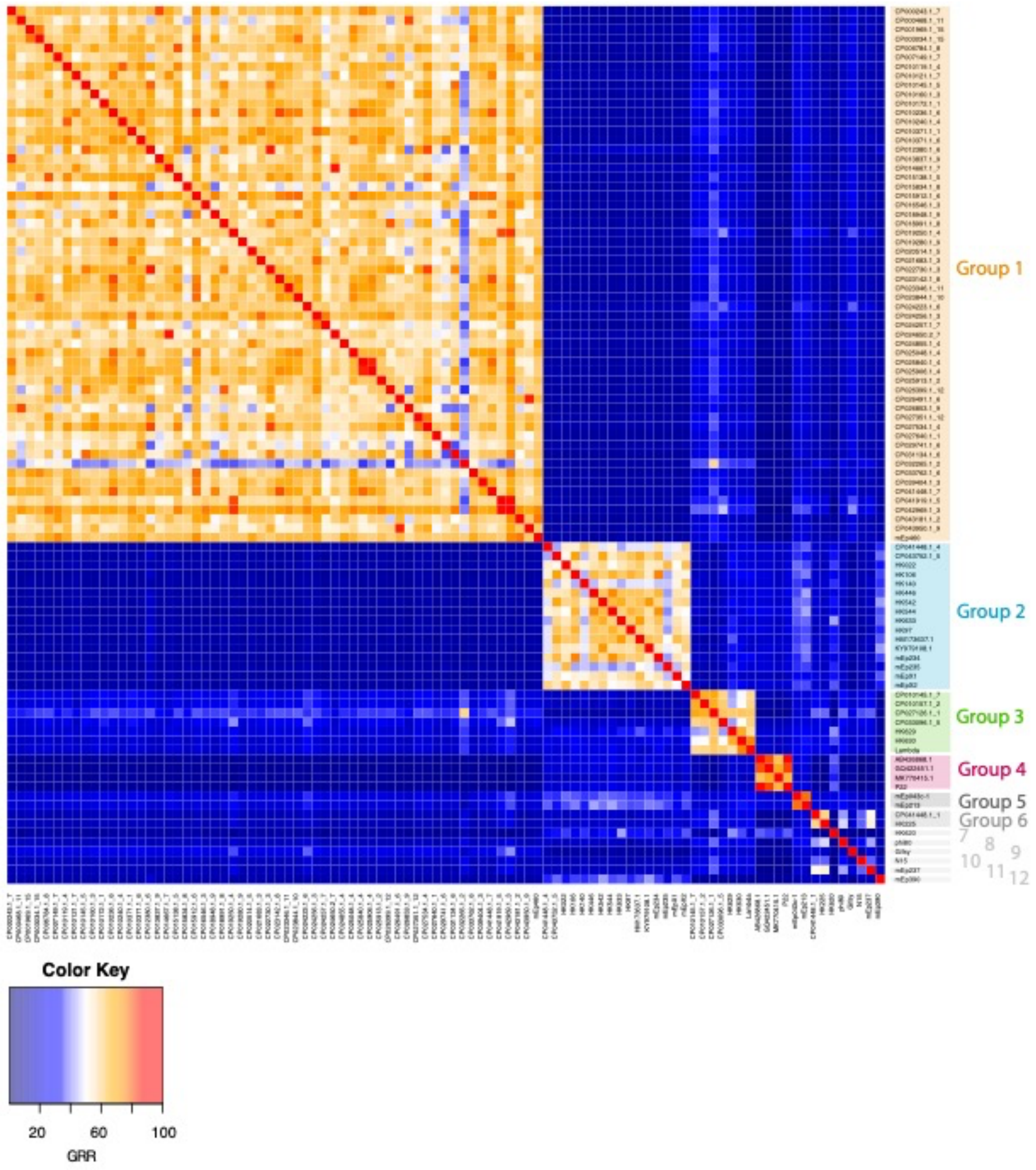


**Figure S2: GRR heatmap for 95 lambdoid phage genomes.** The genome groups are highlighted by colors, where groups with less than 5 members are gray. Groups are sorted by size and the phages within each group are sorted alphabetically (case sensitive).

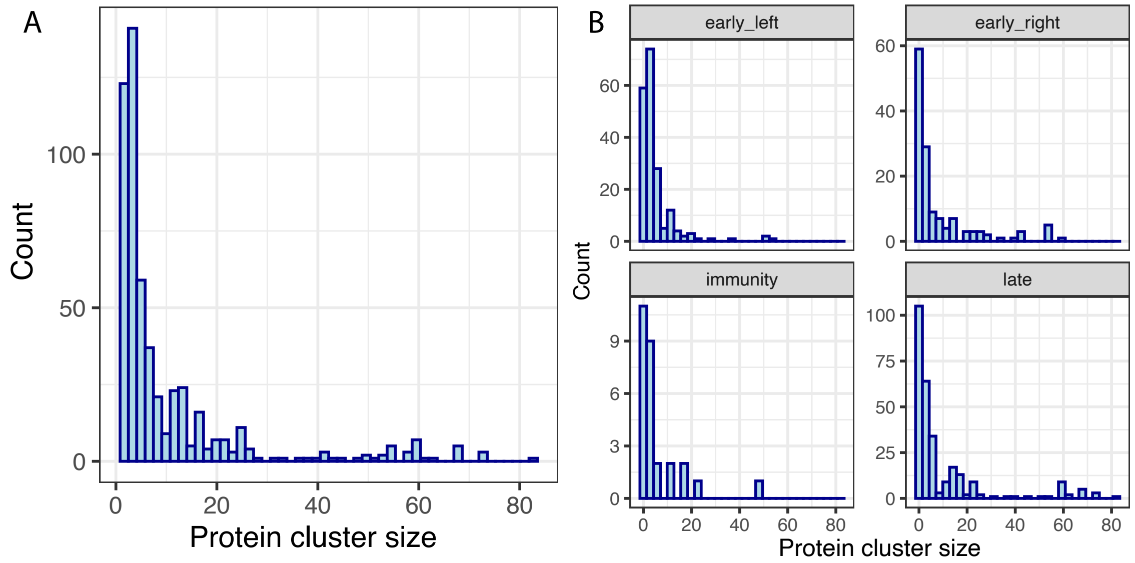


**Figure S3: Protein cluster size distributions.** A) Total protein cluster size distribution, singletons are excluded. B) Protein cluster size distribution per operon, singletons are included, proteins encoded in unknown operons are not shown.

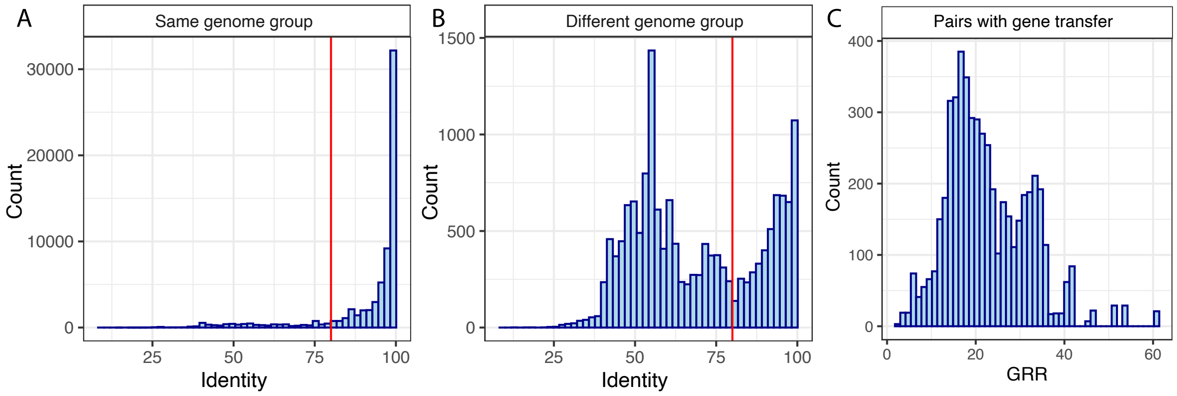


**Figure S4: Cutoff for HGT detection.** A,B) Histogram over pairwise protein identities within protein clusters from A) the same genome group or B) different genome groups. For the same genome group, we observe that 88% of the protein pairs in protein clusters have high identities (above 80%, red line). In contrast, only 5,238 (33%) of the protein pairs from different genome groups have identities above 80%. This motivates the identity cutoff of 80% that is used to detect transferred proteins. C) Histogram over GRR for each transferred protein. Note that a genome pair can be included multiple times if multiple proteins have been transferred.

A

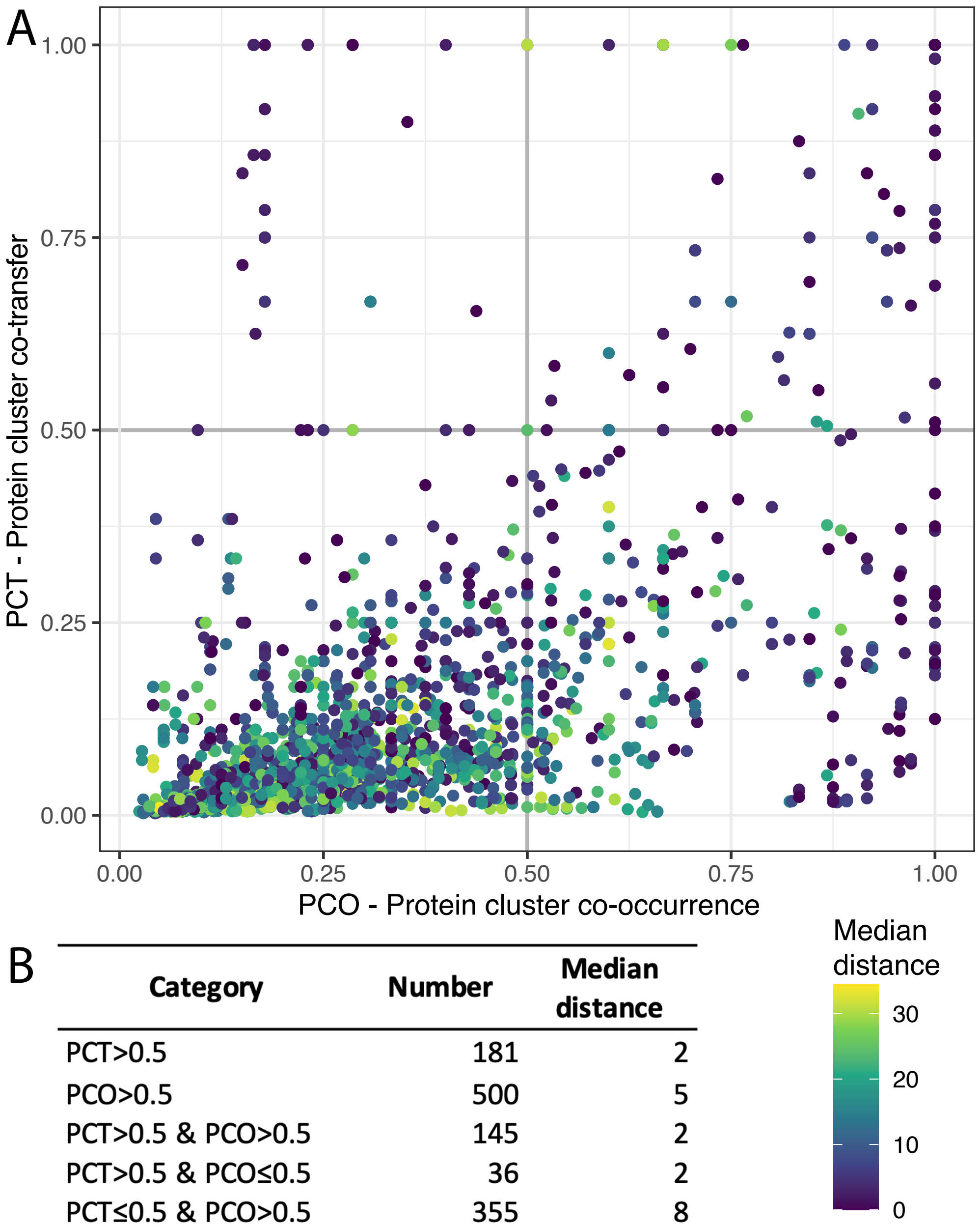


B

| Category | Number | Median distance |
|---|---|---|
| PCT>0.5 | 181 | 2 |
| PCO>0.5 | 500 | 5 |
| PCT>0.5 & PCO>0.5 | 145 | 2 |
| PCT>0.5 & PCO≤0.5 | 36 | 2 |
| PCT≤0.5 & PCO>0.5 | 355 | 8 |

**Figure S5: Protein cluster co-transfer (PCT) and protein cluster co-occurrence (PCO) for all protein clusters that are involved in co-transfer.** A) Color-coded by median distance in number of genes, where the median is calculated across all genomes that contain both protein clusters. B) Number of frequent co-transfers (PCT>0.5), frequent co-occurrence (PCO>0.5) and combinations of these conditions. In addition, the median over all median distances in the category is listed.

**Supplementary tables** (in separate xlsx)

Table S1: Phage genomes included in the analysis. A) Focus phages. B) NCBI phages. C) *E. coli* prophages

Table S2: Genome annotation

Table S3: Pairs of transferred genes

Table S4: Number of transferred protein clusters per operon

Table S5: Pairs of protein clusters that are frequently co-transferred

Table S6: Modules of protein clusters involved in frequent co-transfers

Table S7: Pairwise similarities for Cro, CI, and Kil protein clusters. A) Within-cluster similarities. The proteins marked by a star (*) are involved in recombination. B) Between-cluster similarities. Only the pairs with a significant blast hit are considered for the average identity calculation.
